# Alterations to the cardiac metabolome induced by chronic *T. cruzi* infection relate to the degree of cardiac pathology

**DOI:** 10.1101/2020.09.17.300608

**Authors:** Kristyn Hoffman, Zongyuan Liu, Ekram Hossain, Maria Elena Bottazzi, Peter J. Hotez, Kathryn M. Jones, Laura-Isobel McCall

**Affiliations:** Department of Molecular Virology and Microbiology, Baylor College of Medicine, Houston, TX, United States of America; Department of Pediatrics, Section of Tropical Medicine, Baylor College of Medicine, Houston, TX, United States of America; Department of Chemistry and Biochemistry, University of Oklahoma, Norman, Oklahoma, United States of America; Laboratories of Molecular Anthropology and Microbiome Research, University of Oklahoma, Norman, Oklahoma, United States of America; Texas Children’s Hospital Center for Vaccine Development, Houston, TX; Department of Biology, Baylor University, Waco, TX, United States of America; Department of Microbiology and Plant Biology, University of Oklahoma, Norman, Oklahoma, United States of America

**Author notes:** LIM and KMJ contributed equally to this manuscript. For correspondence (LIM), (KMJ).

**Keywords:** Chagas disease, chronic Chagasic cardiomyopathy, cardiac metabolome

## Abstract

Chronic Chagasic cardiomyopathy (CCC) is a Neglected Tropical Disease caused by the parasite *Trypanosoma cruzi*. The pathognomonic findings in symptomatic CCC patients and animal models includes diffuse cardiac fibrosis and inflammation with persistent parasite presence in the heart. This study investigated chemical alterations in different regions of the heart in relation to cardiac pathology indicators to better understand the long-term pathogenesis of this neglected disease. We used data from echocardiography, fibrosis biomarkers, and histopathological analysis to fully evaluate cardiac pathology. Metabolites isolated from the pericardial and endocardial sides of the right ventricular myocardium were analyzed by liquid chromatography tandem mass spectrometry. The endocardial sections contained significantly less cardiac inflammation and fibrosis than the pericardial sections. Cardiac levels of acylcarnitines, phosphocholines, and other metabolites were significantly disrupted in accordance with cardiac fibrosis, inflammation, and serum fibrosis biomarker levels. These findings have potential implications in treatment and monitoring for CCC patients.

---

Chronic Chagasic cardiomyopathy (CCC) is a neglected tropical disease afflicting over 7 million people worldwide, caused by chronic infection with the parasite *Trypanosoma cruzi*.^1^ The cardiac pathology typically seen in CCC is characterized by an insidious process of cardiac damage from the parasite and host inflammatory response, followed by pathologic remodeling that results in accumulation of cardiac fibrosis.^2–7^ Over 10-15 years after initial infection, the progressive effects of the cardiac inflammation and fibrosis can manifest as multiple cardiac abnormalities, including heart failure, thromboembolism, and/or arrhythmias, any of which may be fatal.^2,8^ Dysfunction of the right ventricle in patients is a strong indicator of poor clinical outcome, including congestive heart failure development after right ventricular dilatation occurs following progressive fibrosis.^9,10^ Current treatment for patients with CCC is restricted to general heart failure and antiarrhythmic medications that are not specific to *T. cruzi* infection. The anti-parasitic treatments available for infected patients, including benznidazole and nifurtimox, have not been found to improve clinical outcome at the advanced symptomatic chronic phase of disease, likely because these drugs do not address the underlying pathologic host response leading to cardiac dysfunction and fail to prevent death in patients with established cardiac disease..^11,12^ There is a dire need for effective drugs to specifically treat the progression of CCC, which require a comprehensive understanding of the stages of cardiac pathogenesis for proper development and implementation.

Previous studies have uncovered some of the cellular signaling that contributes to the development of the cardiac pathology in chronic infection, pointing to both parasite-driven and host-driven signaling mechanisms.^13–19^ Inflammatory pathways driven by interleukin 6, interferon-γ, and tumor necrosis factor-α are important for host control of parasite levels, but also contribute to damage to the myocardium and manifestation of persistent myocarditis.^17,18,20,21^ The pathologic repair mechanism for the damage caused by this inflammation and by the parasites is cardiac fibrosis, controlled by various growth factors, including tissue growth factor-β (TGF-β), connective tissue growth factor (CTGF), and platelet-derived growth factor-D (PDGF-D).^22–25^ These findings illustrated the many changes that occur in the heart throughout CCC, providing some potential targets in inflammatory and fibrotic pathways for therapeutics and vaccines.^22,26–29^ However, further work must be done to uncover the higher resolution details of the nature of changes in the cellular activity of the heart in CCC, including changes to cardiac metabolism, as it is important to understand for functional analysis and processing of drugs that may be effective at preventing or treating cardiac damage.

*T. cruzi* infection has been found to cause perturbations in the tissue metabolome, in a site-specific manner.^30–32,33^ Chemical mapping of the Chagasic heart revealed a differential metabolite profile between heart base and apex, as well as increased parasite numbers at the base of the heart, suggesting a connection between parasite burden and cardiac metabolism.^30^ Additionally, specific acylcarnitine and phosphocholine metabolite levels were found to be significantly correlated with acute Chagas disease severity and mortality in mouse models.^30^ To create a higher resolution understanding of the effects of the cardiac damage by the parasite and the immune response, as well as subsequent repair mechanisms, on the cardiac metabolome, we investigated the metabolic profile of chronically-infected heart tissue with comparisons to cardiac pathology indicators. We hypothesized that the cardiac metabolome could help identify potential biomarkers and represents a source for diagnostic and therapeutic targets that may be effective against cardiac damage in CCC, particularly those closely related to worsening cardiac pathology indicators. To expand upon previous spatial metabolomics findings,^30,33,32^ the current study investigated differences in the cardiac metabolome based on myocardial depth, comparing endocardial-side versus pericardial-side sections of myocardium in infected and naïve mouse models of CCC. In addition, the current study incorporated cardiac pathology indicators, namely cardiac inflammation and fibrosis measured on endpoint histopathology, and circulating biomarkers of cardiac fibrosis, into the metabolome analysis. Machine learning was employed to evaluate the metabolic differences between differing levels of cardiac pathology.

Results indicated that the degree of cardiac pathologies, including cardiac fibrosis and inflammation, correlate with local metabolism of phosphocholines and acylcarnitines. The current findings provide insight into the importance of myocardial depth and degree of cardiac pathology to metabolic pathway disturbances in the heart in CCC, which may have major implications to the efficacy of preventive or therapeutic drugs, as metabolism and distribution in the target tissue may be altered by disease status.

## Results

### Infection and Pathology

In Hoffman et al., 2019, we showed significant elevations in serum levels of TGF-β, CTGF, and PDGF compared to naïve control mice at 209 DPI. Infected groups had an overall mortality of 40% compared with 0% in uninfected age-matched controls.^22^ Chronically infected animals exhibited an overall significant increase in both average cardiac inflammation (p<0.05) and fibrosis (p<0.01) compared to naive control animals at study endpoint (Figure S1).^22^ However, that prior work did not investigate spatial aspects of histopathology.

The current study sought to evaluate cardiac pathology based on myocardial depth in these same animals, producing the following new results. Each individual animal’s cardiac fibrosis and inflammation was significantly increased (p<0.0001, p<0.01, respectively) on the portion of myocardium nearer to the pericardium, compared to the portion of myocardium nearer to the endocardium (Figure 1).

**Figure 1.**
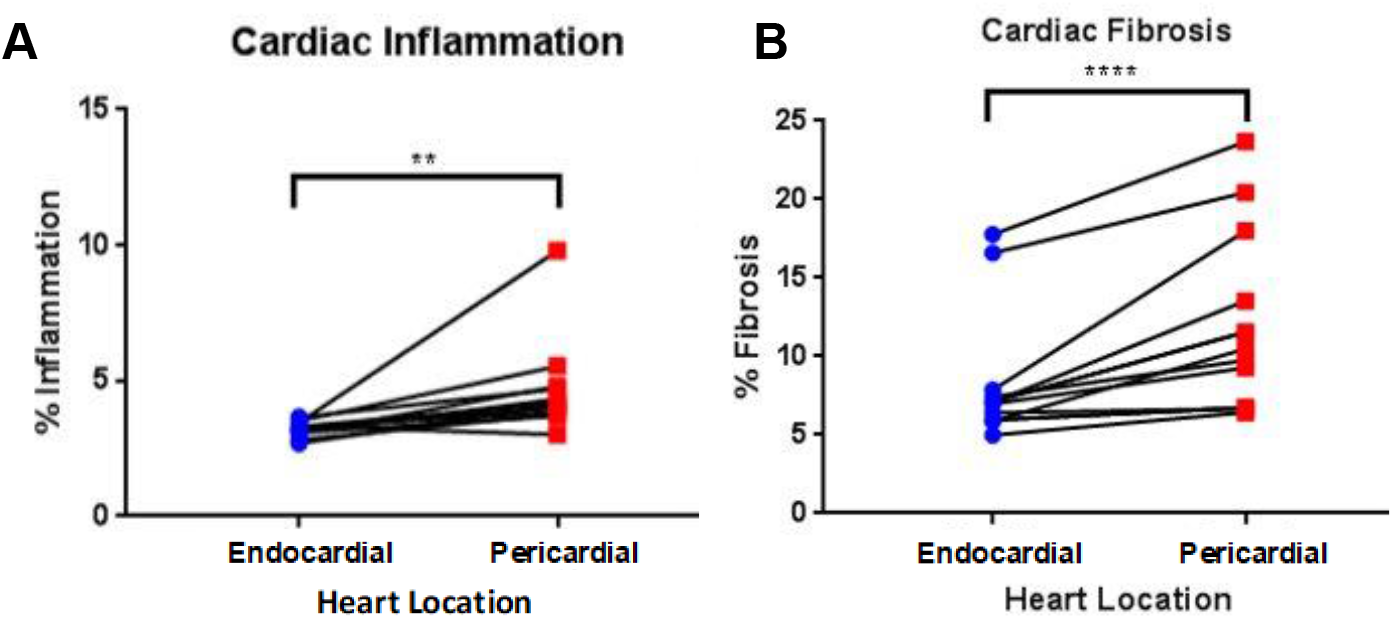
Spatial distribution of fibrosis and inflammation in the infected heart based on myocardial depth (endocardial side, pericardial side). Cardiac inflammation and fibrosis was measured as H&E or Masson’s Trichrome staining, respectively, percent area of total myocardial tissue. Cardiac inflammation (A) was significantly higher on the pericardial aspect of the myocardium of each infected animal. Cardiac fibrosis (B) was also significantly higher on the pericardial aspect of the myocardium of each animal. ** p<0.01, **** p<0.0001

### PCoA analysis reveals pathology-based alterations to the metabolome

Given previous reports of acute *T. cruzi* infection affecting cardiac metabolism, ^30^ it was necessary to investigate whether these changes were also observed in chronic infection, and whether they were correlated to the severity of disease (as quantified by the circulating biomarkers of cardiac fibrosis that we previously reported).^22^ Thus, we performed metabolomic analysis of the right ventricle, divided between endocardial and pericardial segments, in the same animals as above and as in Hoffman *et al*.^22^ As expected, infection status significantly affected the cardiac metabolome (endocardial and pericardial sections combined; PERMANOVA p=0.013, R^2^=12.7%). The overall cardiac metabolome was also significantly affected in relation to indicators of cardiac pathology, including cardiac fibrosis (PERMANOVA p=0.005, R^2^=19.7%), inflammation (PERMANOVA p=0.014 R^2^=12.4%), cardiac fibrosis biomarker concentrations, and circumferential strain measured at endpoint echocardiography (PERMANOVA p=0.006 R^2^=15.0%) (Figure 2 A-F). Of the cardiac fibrosis biomarkers, concentrations of CTGF (PERMANOVA p=0.001 R^2^=22.0%) and TGF-β (PERMANOVA p=0.002 R^2^=19.3%) exhibit the greatest relationship to the cardiac metabolome compared to PDGF concentrations, which did not have a significant relationship with overall metabolome composition (PERMANOVA p=0.12 R^2^=6.25%) (Figure 2C-E). Binning PDGF concentrations into low, medium and high categories did however results in a significant association between cardiac metabolome and this PDGF score (PERMANOVA p=0.04 R^2^=18.9%). No significant differences in cardiac metabolome composition were observed based on myocardial depth by PCoA analysis (PERMANOVA p=0.273 R^2^=3.43%; Figure 2G). However, in accordance with the histological analysis showing worse disease in the pericardial segment (Figure 1), a significant relationship between inflammation score and cardiac metabolome perturbations was only observed for the pericardial cardiac segment (PERMANOVA p=0.043 R^2^=23.4% for infected pericardial samples; non-significant for infected endocardial samples, PERMANOVA p=0.571 R^2^=5.98%).

**Figure 2.**
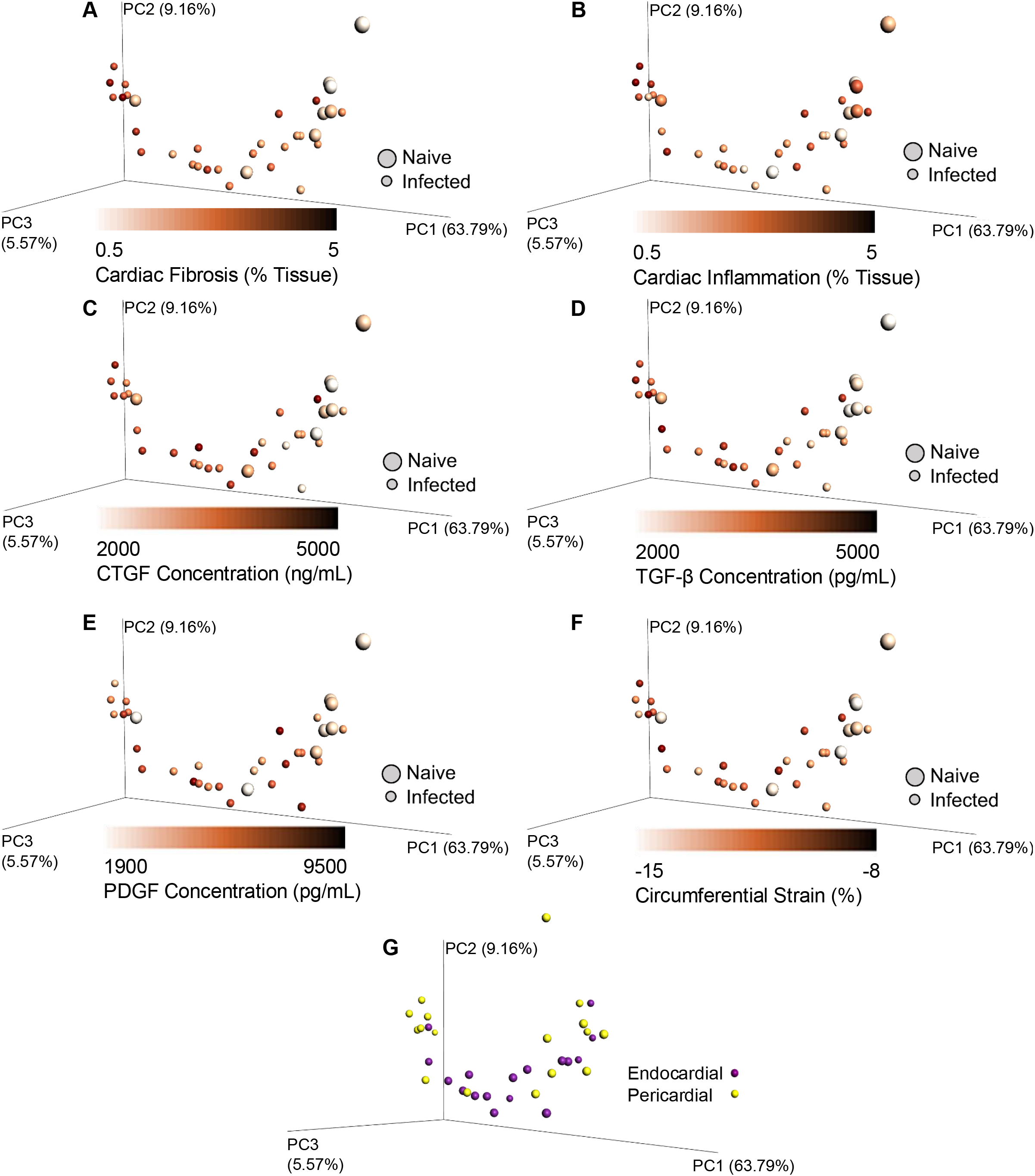
Principal coordinates analysis (PCoA) of dataset by myocardial depth and cardiac pathology level. PCoA plots show a gradient from low and moderate cardiac fibrosis to severe fibrosis (A), low cardiac inflammation to mild inflammation (B), low levels of CTGF (C), TGF-β (D), and PDGF (E) to high levels, increased cardiac strain compared to normal cardiac strain (F). Limited differences were observed between endocardial heart section samples compared to pericardial heart section samples by PCoA (G).

### Relationship between cardiac pathology and cardiac metabolome

We then sought to identify the specific metabolites driving the correlation between metabolomics data and cardiac pathology indicators (Figure 2). Metabolomics data was analyzed by random forest regression based on cardiac fibrosis or inflammation levels and serum levels of CTGF, TGF-β, and PDGF, to identify metabolites that related strongly to indicators of cardiac disease. The metabolites that were highest ranked by random forest were annotated with GNPS.^34,35^ While most metabolites could not be annotated (Table S2), several annotatable metabolites were of physiological significance. Of the annotated metabolites, mainly acylcarnitines and phosphocholines were correlated to serum biomarkers of cardiac fibrosis. All detected metabolites from the acylcarnitine and phosphocholine families in the heart were therefore identified using molecular networking. ^34,35^ The relationships between abundances of all of these metabolites and cardiac fibrosis and inflammation measured on histological samples, as well as serum levels of cardiac fibrosis biomarkers are described below.

### Phosphocholines

Metabolites in the phosphocholine family as identified by molecular networking (Figure 3) and/or found in the top differential features identified by random forest analysis were found to be correlated to several indicators of cardiac pathology. Linear regression analysis revealed positive relationships between LPC(16:0) (*m/z* 496.339, RT 2.99), LPC(18:1) (*m/z* 522.355, RT 2.87), and/or C24:1 Sphingomyelin (*m/z* 813.682, RT 3.33) and cardiac inflammation, cardiac fibrosis, serum TGF-β, and serum CTGF (Table 1). A negative relationship was found between PC(o36:5) (*m/z* 766.572, RT 5.41) and PC(o38:6) (*m/z* 792.587, RT 5.53) versus cardiac fibrosis, serum TGF-β, and serum CTGF (Table 1). When grouped by *m/z* in increments of 100, total phosphocholine metabolites with *m/z* values ranging from 400-500 positively correlated to inflammation, fibrosis, CTGF, and TGF-β levels, those ranging from 500-600 positively correlated to fibrosis, CTGF, TGF-β, and PDGF levels, and those from 800-900 negatively correlated to PDGF levels. When summed, all annotated phosphocholine levels positively correlated to cardiac inflammation serum CTGF levels (Table 2).

**Table 1.** Regression relationships of phosphocholine metabolites and indicators of cardiac pathology in a mouse model of CCC.

| *m/z* | RT (min) | Annotation | Cosine Score | Shared Peaks | Ppm Error | Correlated Pathology Indicator | Linear Regression P values | Linear Regression R<sup>2</sup> values | Linear Regression Slope |
| --- | --- | --- | --- | --- | --- | --- | --- | --- | --- |
| 496.339 | 2.99 | LPC(16:0) | 0.98 | 12 | 1 | Inflammation<br>Fibrosis<br>CTGF<br>TGF-β | 0.0277<br>0.0009<br>0.00004<br>0.0013 | 0.152<br>0.271<br>0.396<br>0.256 | 7.759E-5<br>2.338E-5<br>2.098E-7<br>1.727E-7 |
| 522.355 | 2.87 | LPC(18:1) | 0.93 | 9 | 0 | Inflammation<br>Fibrosis<br>CTGF<br>TGF-β | 0.0062<br>0.0008<br>0.00004<br>0.0012 | 0.187<br>0.274<br>0.396<br>0.258 | 1.587E-5<br>4.338E-6<br>3.873E-8<br>3.197E-8 |
| 766.572 | 5.41 | PC(o36:5) | 0.89 | 8 | 0 | TGF-β | 0.0271 | 0.117 | -5.213E-9 |
| 792.587 | 5.53 | PC(o38:6) | 0.90 | 9 | 3 | Fibrosis<br>CTGF | 0.0341<br>0.01296 | 0.106<br>0.152 | -4.071E-7<br>-3.555E-8 |
| 813.682 | 3.33 | C24:1 Sphingomyelin | 0.94 | 8 | 2 | Inflammation | 0.0057 | 0.191 | 4.760E-5 |

**Table 2.** Regression relationships of phosphocholine metabolite abundances and indicators of cardiac pathology in a mouse model of CCC. No significant relationships with total PCs in the *m/z* range of 200-299, 300-399, 600-699 and 700-799 were observed.

| *m/z* range | Correlated Pathology Indicator | Linear Regression P values | Linear Regression R<sup>2</sup> values | Linear Regression Slope |
| --- | --- | --- | --- | --- |
| 400-499 | Inflammation<br>Fibrosis<br>CTGF<br>TGF-β | 0.005027<br>0.0006406<br>0.00004029<br>0.001163 | 0.1966<br>0.2875<br>0.3959<br>0.2621 | 2.507E-4<br>6.846E-5<br>5.980E-7<br>4.970E-7 |
| 500-599 | Fibrosis<br>CTGF<br>TGF-β<br>PDGF | 0.001726<br>0.00001508<br>0.001204<br>0.03306 | 0.2448<br>0.4306<br>0.2606<br>0.1072 | 2.912E-5<br>2.840E-7<br>2.268E-7<br>5.686E-8 |
| 800-899 | PDGF | 0.03985 | 0.09814 | -1.490E-7 |
| All | Inflammation<br>CTGF | 0.01274<br>0.001449 | 0.1531<br>0.2525 | 4.153E-4<br>8.978E-7 |

**Figure 3.**
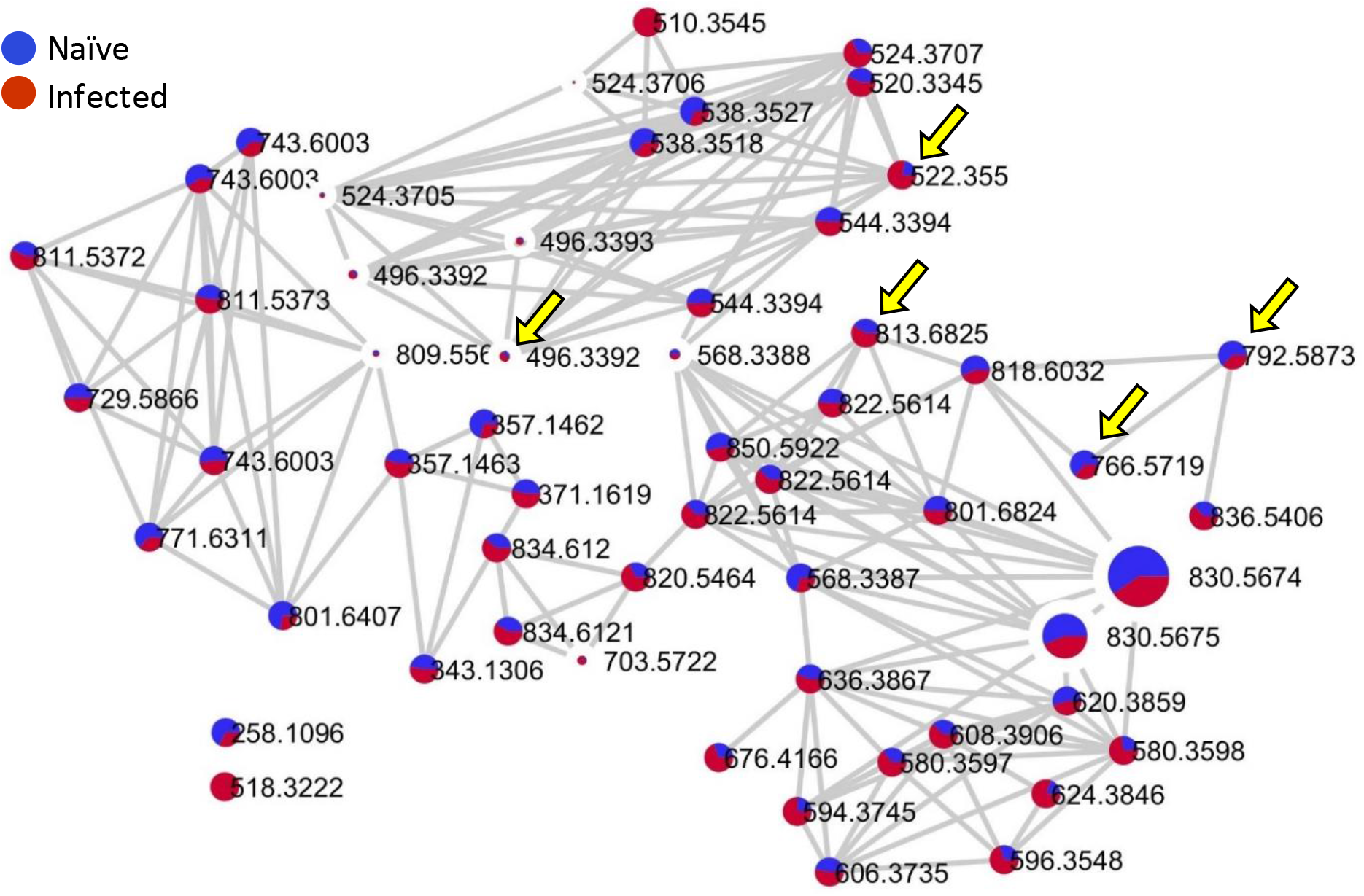
Phosphocholine metabolite family in heart tissue from mice with CCC and naïve control mice. The size of nodes represents relative abundance of each metabolite in the cardiac tissue. All metabolites are labeled with their respective *m/z* values. Yellow arrows indicate the phosphocholine metabolites that were found to have a significant correlative relationship with cardiac pathology indicators (Table 1).

### Alteration of acylcarnitine metabolism based on cardiac pathology

Acylcarnitine metabolites were also found in the top 50 differential features identified by random forest analysis to be related to several indicators of cardiac pathology and thus we investigated the relationship between acylcarnitines and cardiac pathology indicators in more detail (Figure 4). Linear regression analysis revealed a positive relationship between acetylcarnitine (C2:0 acylcarnitine, *m/z* 204.123, RT 0.31), C10:0-OH acylcarnitine (*m/z* 332.242, RT 2.49), and/or C16:0-OH acylcarnitine (*m/z* 416.336, RT 2.53), versus cardiac inflammation, cardiac fibrosis, and/or serum CTGF (Table 3). Additionally, a negative regression relationship was detected between C5:0-OH acylcarnitine (*m/z* 262.164, RT 0.73) versus serum TGF-β and CTGF. On a global level, mid-chain (C5 to C11) acylcarnitines were negatively correlated with PDGF.

**Figure 4.**
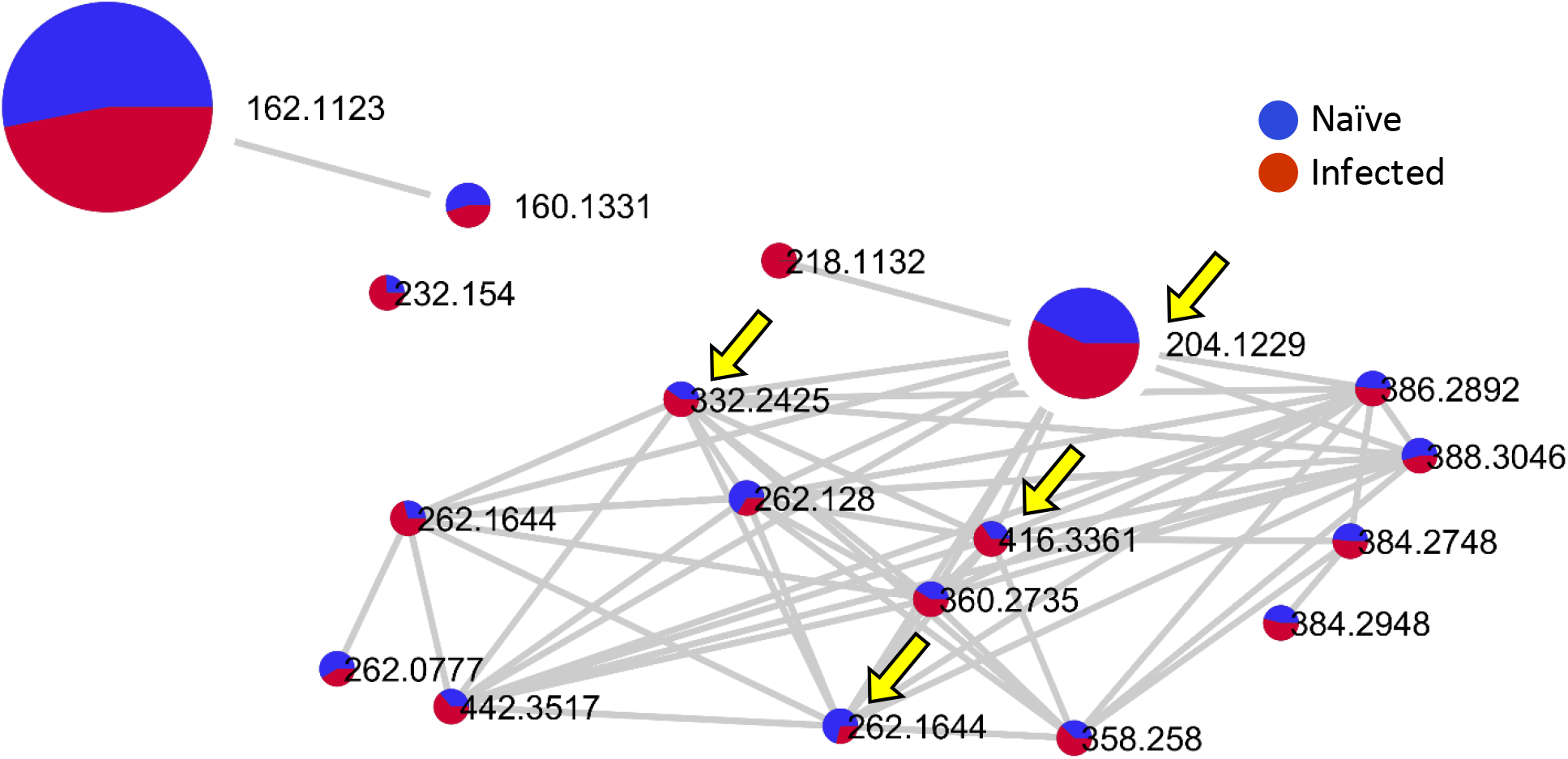
Acylcarnitine metabolite family in heart tissue from mice with CCC. The size of nodes represents relative abundance of each metabolite in the cardiac tissue. All metabolites are labeled with their respective *m/z* values. Yellow arrows indicate the three acylcarnitine metabolites that were found to have a significant correlative relationship with cardiac pathology indicators (Table 3).

**Table 3.** Regression relationships of acylcarnitine metabolites and indicators of cardiac pathology in a mouse model of CCC.

| *m/z* | RT (min) | Annotation | Cosine Score | Shared Peaks | Ppm Error | Correlated Pathology Indicator | P values | R<sup>2</sup> values | Slope |
| --- | --- | --- | --- | --- | --- | --- | --- | --- | --- |
| 204.123 | 0.31 | C2:0 acylcarnitine | 0.98 | 6 | 0 | Inflammation<br>Fibrosis | 0.0002<br>0.0137 | 0.337<br>0.150 | 2.176E-4<br>3.521E-5 |
| 262.164 | 0.73 | C5:0-OH acylcarnitine | 0.98 | 9 | 1 | TGF-β<br>CTGF | 0.0235<br>0.0054 | 0.124<br>0.193 | -1.038E-8<br>-1.244E-8 |
| 332.242 | 2.48 | C10:0-OH acylcarnitine | 0.8 | 9 | 0 | CTGF | 0.0088 | 0.170 | 1.592E-8 |
| 416.336 | 2.90 | C16:0-OH acylcarnitine | 0.93 | 9 | 2 | CTGF | 0.0424 | 0.095 | 1.180E-8 |
| Mid-chain acylcarnitines (C5-C11) |  |  | NA | NA | NA | PDGF | 0.0361 | 0.103 | -3.492E-9 |

### Physiologic consequences of infection: drug metabolism

Strikingly, several members of the ketamine metabolite network were also found in the top random forest differential features correlated to several indicators of cardiac pathology. Ketamine was administered to all animals as a component of the anesthetic cocktail for endpoint tissue collection. As the dosing schedule (150 mg/kg body weight administered at euthanasia) and amount of ketamine administered were the same for all animals, the differences in ketamine metabolite abundances in the heart are due to infection status and cardiac pathology severity. Ketamine (*m/z* 207.057, RT 2.33) was in significantly greater abundance (p<0.01) in the pericardial section of the heart than the endocardial section, suggesting an alteration in the positional distribution, metabolism, and/or clearance of ketamine in the heart (Figure 5B). These findings may reflect positional changes in the cardiac pathology during chronic infection, including increased inflammation and fibrosis in the pericardial section of the heart compared to the endocardial section (Figure 1). In addition, the total sum of all ketamine metabolite abundances in the heart tissue was significantly greater (p<0.01) in infected mice compared to naïve mice (Figure 5C). Regression analysis revealed a positive relationship between ketamine, norketamine, and two desmethylketamine isomers versus cardiac inflammation, cardiac fibrosis, serum TGF-β, and/or serum CTGF (Table 4).

**Figure 5.**
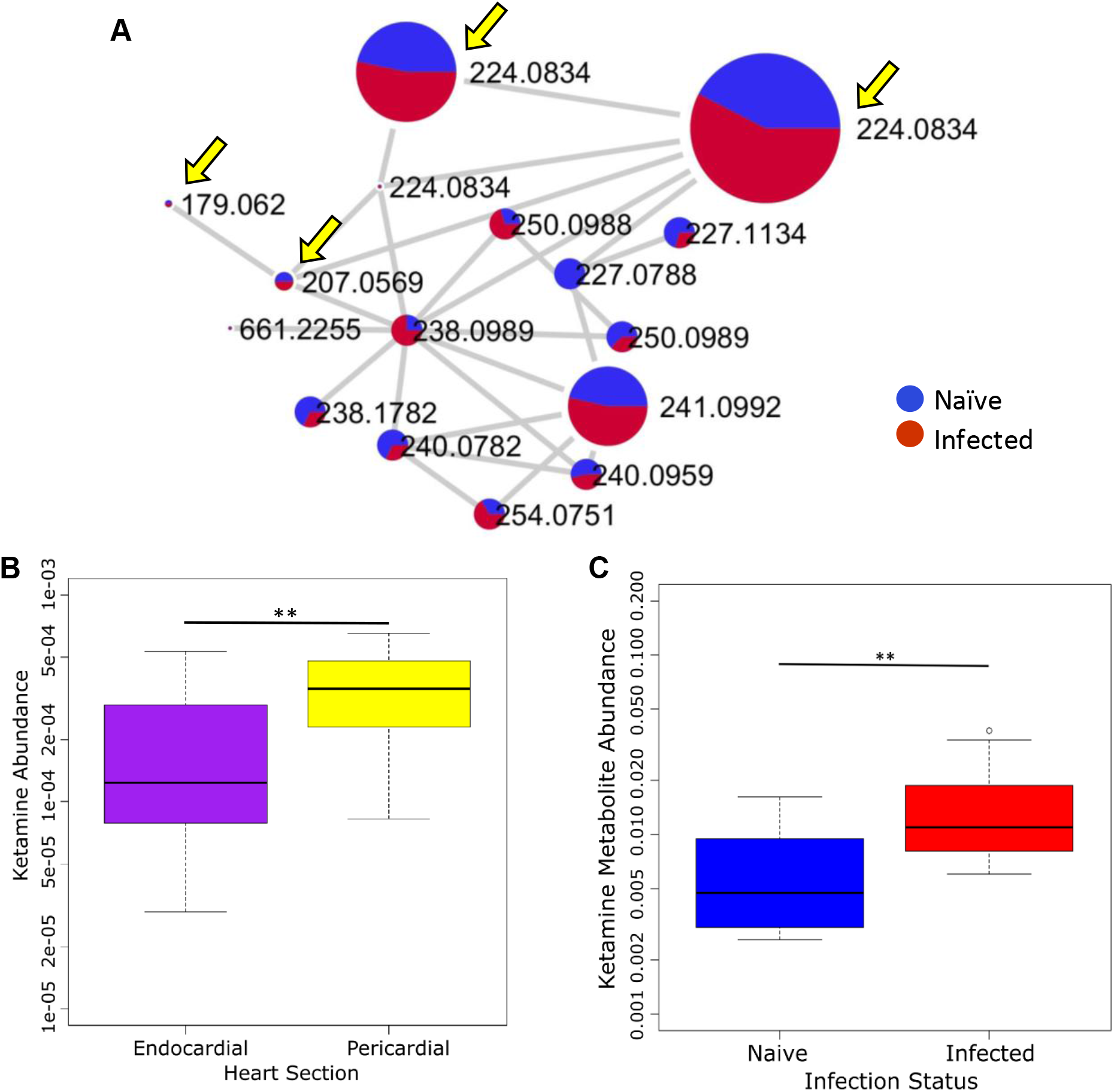
Ketamine metabolites by infection status and myocardial depth. Ketamine metabolite family in heart tissue from mice with CCC (A). The size of nodes represents relative abundance of each metabolite. All metabolites are labeled with their respective *m/z* values. Yellow arrows indicate the ketamine metabolites that were found to have a significant correlative relationship with cardiac pathology indicators (Table 4). Ketamine (B) metabolite abundances were significantly higher in the pericardial versus endocardial myocardial section of all mice. Total ketamine metabolite abundance was significantly higher in infected versus naïve mice (C). ** p<0.01, *p<0.05.

**Table 4.**
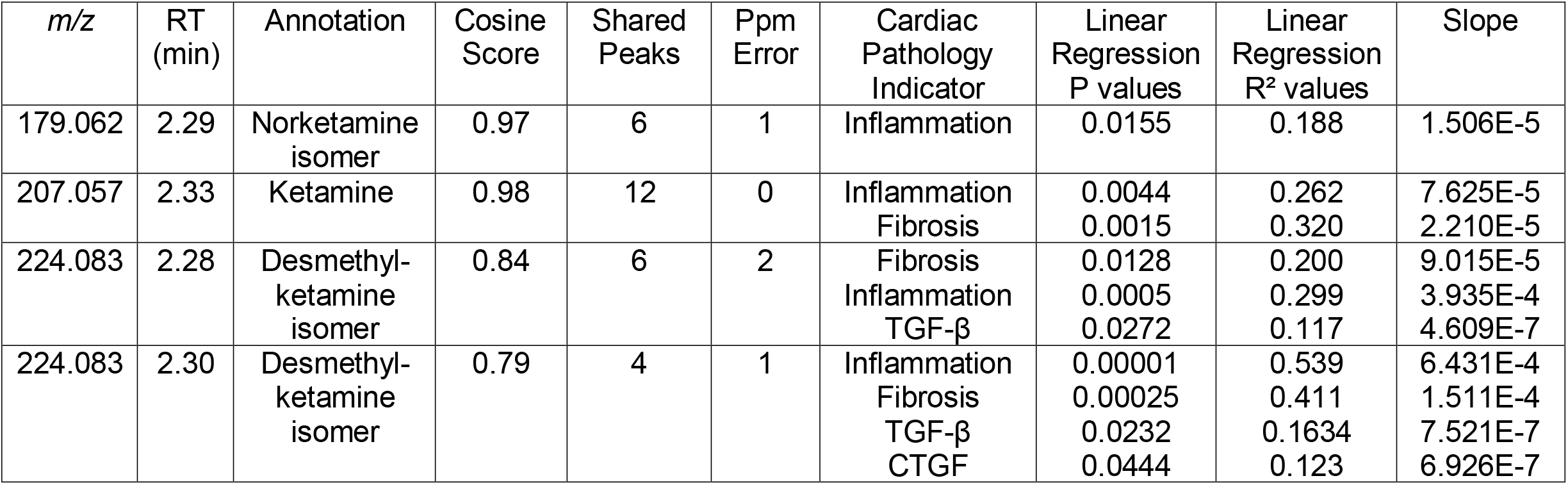
Regression relationships of ketamine metabolites versus cardiac pathology indicators in a mouse model of CCC.

| *m/z* | RT (min) | Annotation | Cosine Score | Shared Peaks | Ppm Error | Cardiac Pathology Indicator | Linear Regression P values | Linear Regression R<sup>2</sup> values | Slope |
| --- | --- | --- | --- | --- | --- | --- | --- | --- | --- |
| 179.062 | 2.29 | Norketamine isomer | 0.97 | 6 | 1 | Inflammation | 0.0155 | 0.188 | 1.506E-5 |
| 207.057 | 2.33 | Ketamine | 0.98 | 12 | 0 | Inflammation<br>Fibrosis | 0.0044<br>0.0015 | 0.262<br>0.320 | 7.625E-5<br>2.210E-5 |
| 224.083 | 2.28 | Desmethyl-ketamine isomer | 0.84 | 6 | 2 | Fibrosis<br>Inflammation<br>TGF-β | 0.0128<br>0.0005<br>0.0272 | 0.200<br>0.299<br>0.117 | 9.015E-5<br>3.935E-4<br>4.609E-7 |
| 224.083 | 2.30 | Desmethyl-ketamine isomer | 0.79 | 4 | 1 | Inflammation<br>Fibrosis<br>TGF-β<br>CTGF | 0.00001<br>0.00025<br>0.0232<br>0.0444 | 0.539<br>0.411<br>0.1634<br>0.123 | 6.431E-4<br>1.511E-4<br>7.521E-7<br>6.926E-7 |

## Discussion

Host response to chronic *T. cruzi* infection is a complex, multifactorial process that likely includes many targets for potentially effective therapeutics against development and/or progression of CCC. In this study, we found that the cardiac metabolome of the right ventricle was perturbed based on the degree of chronic CCC pathology, supplementing our prior findings that cardiac metabolome differed between infected and uninfected animals.^30,32^ Cardiac inflammation and fibrosis severity, myocardial strain, and levels of fibrosis biomarkers correlated with changes in the metabolism of phosphocholines and acylcarnitines (Figure 2, Tables 2-3).

We and others previously identified perturbation in acylcarnitine metabolism induced by *T. cruzi* infection when comparing infected vs uninfected animals.^30–33,36^ In the current study, we found alterations in cardiac tissue acylcarnitine metabolite abundances correlate to severity of cardiac inflammation and fibrosis, as well as serum levels of cardiac fibrosis biomarkers, with mid-chain acylcarnitines in particular negatively correlated with serum PDGF. These results complement previous findings and concur with our prior observation of reduced mid-chain acylcarnitines in the heart of C3H/HeJ mice chronically infected with two different *T. cruzi* strains, Sylvio X10/4 (DTU TcI) and CL (DTU TcVI).^32^ One limitation, however, comparing these studies is that no acylcarnitines longer than C16 were retained after data processing and filtering in this study, unlike in prior work.

Overall, these observations reinforce the strength of the evidence supporting acylcarnitines as potential biomarkers of disease and potential therapeutic targets. Additionally, we recently reported findings that carnitine supplementation reduced acute mortality in a mouse model of Chagas disease, mitigated the metabolic disturbances caused by infection in the heart and reduced markers of heart stress (brain natriuretic peptide (BNP)).^33^ The use of acylcarnitines and other metabolites as biomarkers of cardiac pathology severity could also be applied to preclinical studies of therapeutic efficacy, to offer supporting information regarding a drug’s effect on CCC at the metabolic level. This would improve the resolution of preclinical efficacy studies for Chagas disease and represents future directions of these findings.

Phosphocholine metabolism was also found to be affected by severity of infection with *T. cruzi* in this study. *T. cruzi* infection has been previously found to alter levels of members of the phosphocholine metabolic family in the serum and heart.^30,32,36^ Our observation of positive correlation between overall phosphocholines and phosphocholines in mass ranges *m/z* 400-499, 500-599 vs fibrosis and/or inflammation concurs with our observation of infection-induced elevation in phosphocholines in an independent chronic infection system.^32^ Phosphocholines are key factors in metabolism of lipids, choline production, and membrane structure.^37–40^ Metabolism of phosphocholines has been found to be altered in heart disease, including ischemia with reperfusion injury.^41^ It is possible that the hallmarks of CCC pathology, myocarditis and progressive cardiac fibrosis, cause continual cell damage, turnover, and tissue architecture remodeling, which results in alterations to phosphocholine metabolism. Further investigation is needed to identify which metabolic processes are impacted by the alterations seen in phosphocholine metabolite levels. These may represent potential diagnostic or therapeutic targets that could detect or ameliorate the pathologic mechanisms behind the myocarditis and cardiac fibrosis in CCC.

In these experiments, we use a well-established mouse model of CCC with progressive cardiac fibrosis and mild to moderate cardiac inflammation.^42^ We found that the degree of severity of cardiac inflammation and fibrosis varies at different myocardial depths, with the pericardial portion on the myocardium exhibiting greater levels of cardiac inflammation and fibrosis (Figure 1). The current study is the first to formally demonstrate any pattern of inflammation and/or fibrosis formation depending on myocardial depth. The finding that cardiac inflammation and fibrosis are more severe in the pericardial region of the heart, closer to the pericardium than the endocardial region of the left ventricle, suggests that the host’s response to the infection is site-specific. The reasoning behind the differences in cardiac pathology based on myocardial depth may be based on parasite location, blood flow or supply of nutrients to each depth level, preferential recruitment of inflammatory cells and fibroblasts to the pericardial depth segment, or other factors. Perfusion of nutrients and key cellular signaling molecules may differ between the two depths of myocardium studied here; indeed, coronary vasculature directionality flows from pericardial to endocardial regions.^43,44^ In addition, there was higher recruitment of inflammatory cells to the pericardial section of the myocardium (Figure 1). Further studies are needed to reveal the reason(s) behind the differential development of cardiac fibrosis and intensity of cardiac inflammation between heart regions during CCC.

The finding that ketamine metabolism is also related to both myocardial depth and cardiac pathology severity has major clinical applications. Importantly, differences in microvasculature could have large impacts on the rate of clearance for ketamine and other drugs. The bioavailability, metabolism, and clearance of any drug in the different depths of the heart will also have great importance on the efficacy against the parasite or further cardiac pathology development. We have previously published that parasite levels differ between cardiac locations,^30^ which, coupled with the current findings of differences in pathology based on location lead us to believe that future studies with Chagas-specific therapeutics and/or preventatives need to include evaluation of the abundance of these drugs and their metabolites at different locations within the heart. It is important that any drugs targeting *T. cruzi*, like benznidazole, or those targeting pathologic host responses, have optimal concentrations, clearance, and metabolism at the most-affected locations throughout the myocardium. Any anti-fibrotic or anti-inflammatory treatment for CCC would likely need greater concentration at the pericardial section of the myocardium, considering the increased severity of both fibrosis and inflammation in this section.

Using this mouse model of CCC, we have previously produced evidence of cardiac disease, including increased cardiac strain, cardiac inflammation and fibrosis, and elevations in biomarkers of cardiac fibrosis.^22^ While all animal models’ limitations must be considered when translating these results to implications for human CCC, the results from the present study, in combination with previous findings, advance our understanding of the relationship between cardiac pathology severity and the cardiac metabolome. The metabolic disturbances provide greater resolution to the pathophysiologic profile of Chagas disease, particularly as they correlate to findings on the tissue level and in circulation that have been connected to disease severity. Importantly, these findings in the subclinical disease seen in our model can be applied to the prolonged, asymptomatic stage of disease experienced by infected humans before development of symptoms, when diagnostics and treatment initiation are most critical. These findings have major implications for consideration and development of targeted therapeutics that could delay or prevent worsening of cardiac disease due to chronic infection with *T. cruzi*.

## Conclusion

The present study found a connection between metabolomic disturbances and cardiac pathology severity in a mouse model of CCC. Acylcarnitine and phosphocholine metabolite abundances were correlated with markers of cardiac inflammation and fibrosis. Pharmacological agent concentrations in the heart were also correlated to both markers of cardiac disease and myocardial depth. Taken together, these findings suggest that chronic *T. cruzi* infection exhibits biological impacts on the cardiac metabolome, with alterations to drug metabolism in a positional manner. Future applications of these findings can uncover the extent to which these biological impacts affect metabolism of treatment or vaccine candidates for Chagas disease, or uncover metabolic pathways that can serve as targets for potential therapeutics or diagnostics.

## Methods

### Animal model of CCC

This study uses the same animals as reported in Hoffman et al., 2019 ^22^. The right ventricle of the hearts of mice from this previous study were frozen at study endpoint for future analysis. Upon assessment of the findings of this prior study, we designed the current study to investigate the metabolite profiles of these right ventricle samples as they relate to our higher resolution findings regarding cardiac pathology through imaging and serum biomarker concentrations in the same animals. Methods used in this previously-reported animal infection, and new methods regarding the current experiments, are stated below.

As previously reported,^22^ all experiments using animals were completed within compliance with the *National Institutes of Health Guide for the Care and Use of Laboratory Animals, 8^th^ Edition*, under a protocol approved by the Baylor College of Medicine Institutional Animal Care and Use Committee.^45^ Female BALB/c mice, aged 6 to 8 weeks, were divided into two groups and infected with 500 *T. cruzi* H1 (TcI) blood-form trypomastigotes suspended in 0.1 mL sterile saline (n=20) or sterile saline alone (n=4), as previously described.^22,46,47^ Mice were evaluated daily for systemic signs of disease, including ruffled coat, lethargy, hunched posture, dyspnea, and visible weight loss. These signs were used as end point determinants and mice that reach end point were humanely euthanized.

Successful infection was assessed by evaluation of parasitemia, as previously described and reported.^47^ The mice were monitored for 209 DPI and animal survival was monitored over the course of the infection. At study endpoint, all mice were imaged with echocardiography. Following imaging, all mice were euthanized and hearts were collected and sectioned. The right ventricle was frozen (for metabolite analysis) and the left ventricle was immersed in 10% neutral buffered formalin (for histological processing). Serum from the mice was collected at endpoint and immediately frozen. Echocardiography and analysis of images was performed as previously reported and the data from these images were used in the current study, as well.^22^ Serum harvested from endpoint blood collected from all mice was used to measure circulating concentrations of TGF-β, CTGF, and PDGF-D by enzyme-linked immunosorbent assay (ELISA, TGF-β: Thermo Fisher Scientific, Waltham, MA; CTGF and PDGF-D: G-Biosciences, St. Louis, MO; Abclonal, Woburn, MA) as previously reported and the data from these prior experiments was used in the current investigations.^22^ Each experiment was performed with two technical replicates.

### Histopathology of Cardiac Inflammation and Fibrosis

As previously reported, ^22^ endpoint cardiac tissue was removed from euthanized mice, divided into the left and right ventricle sections with a single, long-axis cut from apex to base, and the left ventricle section was immediately fixed in 10% formalin solution for histopathological analysis. The formalin-fixed samples were dehydrated, embedded in paraffin, sectioned on the short axis to 5 μm, fixed, and stained with hematoxylin-eosin and Masson’s trichrome for the quantification of inflammation and fibrosis, respectively.

Analysis of the stained sections of the left ventricle was performed on 3 representative images of the left ventricle, at the basal, apical, and midventricular regions. Images of the sections were obtained with an AmScope ME580 bright field microscope (AmScope, Irvine, CA) equipped with LMPLAN40-065 40X objective using an 18-megapixel camera at fixed upper and lower light levels. These images were divided into endocardial and pericardial (out section halves at 50% ventricular thickness. The images were analyzed using ImageJ software (National Institutes of Health, Bethesda, MD)^48^ to obtain quantitative measurements of inflammation on the hematoxylin-eosin–stained sections and fibrosis on the Masson’s trichrome–stained sections. Briefly, for total fibrosis and inflammation, a numerical value of the total myocardium and the fibrosis or inflammation volume were obtained. The fraction of total volume of fibrosis or inflammation per area of the myocardial section analyzed was calculated for each animal. Total myocardial area was quantified as the area of inflammation or fibrosis added to the area of myocardium (white background in the images was subtracted from total area). Myocardial fibrosis and inflammation values for infected and naive animals were compared with Prism software version 8.0 (GraphPad Software, La Jolla, CA) using the student’s t test, with p values <0.05 considered significant.

### Metabolite extraction

All samples were processed using a method adapted from Want *et al*, as previously implemented in our work.^30,33,49^ The right ventricle samples collected at study endpoint were immediately frozen and stored at −80° Celsius. Right ventricular heart sections (30-50 mg/sample) were thawed and carefully dissected to obtain two equivalent sections of myocardial tissue, which included an endocardial-adjacent section and a pericardial-adjacent section. The heart sections were suspended in LC-MS-grade water (10 μL per 1 mg sample) and homogenized with 5 mm stainless steel beads using a Qiagen TissueLyzer in 2 mL Safelock Eppendorf tubes (3 minutes at 25 Hz). Following homogenization, cold LC-MS-grade methanol with 4 μM sulfachloropyridazine internal standard was added to the homogenate to achieve a final concentration of 50% methanol. All samples were then homogenized again (3 minutes at 25 Hz), then centrifuged at 16,000 G for 10 minutes at 4° Celsius. The supernatant, containing the aqueous extract, was collected and dried in a vacuum concentrator overnight. The pellet was homogenized in 3:1 dichloromethane:methanol, spiked with 2 μM sulfachloropyridazine internal standard (20 μL per 1 mg sample; 6 minutes at 25 Hz). Samples were centrifuged at 16,000 G for 10 minutes at 4° Celsius. The supernatant, containing the organic extract, was collected and air-dried overnight. All samples were then stored at −80° C until LC-MS/MS analysis.

### LC-MS/MS data acquisition

Organic and aqueous heart extracts were resuspended in 50% methanol containing 0.5 μg/mL sulfadimethoxine internal standard and combined. LC-MS/MS was performed as previously described, in randomized sample order.^33^ LC separation was performed on a Thermo Vanquish instrument using a 1.7μm Kinetex C8 50 x 2.1 mm column, 100 Å pore size for right ventricle tissue samples. Chromatography was performed at a flow rate of 0.5 mL/min with water + 0.1% formic acid as mobile phase A and acetonitrile + 0.1% formic acid as mobile phase B. The LC gradient for right ventricle samples was: 0-1 min, 2% B; 1-2.5 min, ramp up linearly to 98% B; 2.5-4.5 min, hold at 98% B; 4.5-5.5 min, ramp down to 2% B; 5.5-7.5 min hold at 2% B. MS/MS analysis was performed on a Q Exactive Plus (ThermoScientific) mass spectrometer. Ions were generated by electrospray ionization and MS spectra acquired in both positive and negative ion mode. Instrumental performance was assessed during the run by monitoring blanks and pooled quality control samples (at start, end, and every 12 injections), and a standard mix of 6 compounds (sulfamethazine, sulfadimethoxine, sulfachloropyridazine, coumarin-314, sulfamethizole, amitriptyline) at start, end and every 100 injections. Full parameters are listed in Table S1.

### LC-MS/MS data analysis

The raw MS datasets were converted to mzXML format with MSconvert and imported into MZmine (version 2.33).^50,51^ Table 5 contains the parameters used to identify MS features.

**Table 5.** Parameters for MZmine data processing.

| MZmine Parameters |  |  |
| --- | --- | --- |
| MS<sup>1</sup> | Retention Time (min.) | 0.0 – 7.51 |
|  | Noise Level | 2.0E6 |
| MS<sup>2</sup> | Retention Time (min.) | 0.0 – 7.51 |
|  | Noise Level | 1.0E3 |
| Chromatogram<br>Builder | Mass List | masses |
|  | Minimum Time Span (min.) | 0.01 |
|  | Minimum Height | 6.0E6 |
|  | *m/z* Tolerance (ppm) | 10.0 |
|  | Algorithm | Baseline Cut-off |
| Chromatogram<br>Deconvolution | *m/z* Range for MS<sup>2</sup> Scan Pairing (Da) | 0.01 |
|  | RT Range for MS<sup>2</sup> Scan Pairing (min.) | 0.2 |
|  | Minimum Peak Height | 6.0E6 |
|  | Peak Duration Range (min.) | 0.01 – 3.50 |
|  | Baseline Level | 2.0E6 |
| Deisotoping | *m/z* Tolerance (ppm) | 10.0 |
|  | Retention Time Tolerance (min.) | 0.5 |
|  | Monotonic Shape | Checked |
|  | Maximum Charge | 3 |
|  | Representative Isotope | Lowest *m/z* |
| Join Alignment | *m/z* Tolerance (ppm) | 10.0 |
|  | Weight for *m/z* | 1 |
|  | Weight for Retention Time | 1 |
|  | Retention Time Tolerance (min.) | 0.5 |
| Feature List<br>Row Filtering | Minimum Peaks in a Row | 6 |
|  | Retention Time (min) | 0.20 – 6.45 |
|  | Keeps Only Peaks with MS<sup>2</sup> Scans | Checked |
|  | Reset Peak No. ID | Checked |

Blanks were removed (3x cutoff) and data was normalized to total ion current (TIC normalized). Principal coordinates analysis (PCoA) was performed on the normalized data using the Bray-Curtis-Faith dissimilarity metric in QIIME1 and were visualized in EMPeror.^52–55^ PERMANOVA analyses were performed in R using the package “vegan.”

Feature-based molecular networks were created using the Global Natural Products Social Molecular Networking platform (GNPS).^34,35^ The parameters for the spectra and library searches were: 0.02 precursor ion mass tolerance and MS/MS fragment ion mass tolerance, ≥0.7 cosine score, ≥4 matched peaks, and 200 Da maximum analog search mass difference. The molecular networks were imported into Cytoscape (v3.7.1) for visualization.^56^

Differential features were identified using random forest analysis of data from infected heart samples in R, with 1,000 trees. The most differential metabolites by random forest analysis and those found in the acylcarnitine, ketamine, and phosphocholine networks created by Cytoscape were annotated according to analog library hits in GNPS and mirror plots visually inspected (Figure S2). Linear regression analysis of the relationship between these metabolites and others in the phosphocholine, acylcarnitine, and ketamine networks defined by Cytoscape and the cardiac pathology indicators, including cardiac inflammation and fibrosis and serum levels of cardiac fibrosis biomarkers was performed in R. Statistical significance of the relationship between ketamine metabolite abundances versus infection status and heart section data was performed using the Wilcoxon rank sum test. Boxplots illustrate the mean, first and third quartile of all samples, with whiskers representing 1.5 times interquartile range.

### Data availability

LC-MS/MS data has been deposited in MassIVE (massive.ucsd.edu), accession number MSV000083798. GNPS analyses can be accessed at: https://gnps.ucsd.edu/ProteoSAFe/status.jsp?task=e8e4466192a34c8db3f711744effd65d.

## Acknowledgements

This work was funded in part by internal funds from the Texas Children’s Hospital Center for Vaccine Development. Equipment and reagents used were funded in part by Baylor Research Advocates for Student Scientists, as well as travel for LC/MS and data analysis training. This project was supported by the Mouse Phenotyping Core at Baylor College of Medicine with funding from the NIH (UM1HG006348) and NIH (RO1DK114356), the Pathology and Histology Core at Baylor College of Medicine with funding from the NIH (P30 CA125123). This project also received partial support from start-up funds from the University of Oklahoma and from the Southwest Electronic Energy Medical Research Institute via a sub-award from Baylor College of Medicine to University of Oklahoma.

