## Supplemental Tables 1,2 for "Alterations to the cardiac metabolome induced by chronic *T. cruzi* infection relate to the degree of cardiac pathology"

**Table S1. Instrumental and LC-MS data processing parameters used.**

| **Instrumental Methods** | | | |
| --- | --- | --- | --- |
| Use Lock Masses | Off | | |
| Chromatogram Peak Width | 6 seconds | | |
| Method Duration | Hearts: 7.5 minutes | | |
| Exclusion List | 371.1010 | | |
|  | 235.2060 | | |
|  | 311.0840 | | |
|  | 314.1310 | | |
|  | 285.0135 | | |
|  | 144.9822 | | |
| **LC Parameters** | | | |
| **Time (minutes)** | **Flow (mL/min)** | **%B (Acetonitirle + 0.1% Formic Acid)** | **Curve** |
| 0.00 | 0.50 | 2.0 | 5 |
| 1.00 | 0.50 | 2.0 | 5 |
| 2.50 | 0.50 | 98.0 | 5 |
| 4.50 | 0.50 | 98.0 | 5 |
| 5.50 | 0.50 | 2.0 | 5 |
| 7.50 | 0.50 | 2.0 | 5 |
| 7.50 | Stop Run | | |
| **MS Parameters** | | | |
| *Source* | | | |
| Sheath gas flow rate | 35 L/min | | |
| Auxiliary gas flow rate | 10 L/min | | |
| Sweep gas flow rate | 0 L/min | | |
| Spray voltage | 3.80 kV | | |
| S-lens RF level | 50 V | | |
| Capillary temperature | 320⁰ C | | |
| Auxiliary gas temperature | 350⁰ C | | |
| *Full MS* | | | |
| Resolution | | 70,000 | |
| AGC Target | | 1E6 | |
| Maximum IT | | 246 milliseconds | |
| Scan Range | | 70 to 1050 *m/z* | |
| *dd-MS^2^* | | | |
| Resolution | | 17,500 | |
| AGC Target | | 2E5 | |
| Maximum IT | | 54 milliseconds | |
| TopN | | 5 | |
| Isolation Window | | 1.0 *m/z* | |
| Fixed First Mass | | -- | |
| (N)CE / Stepped | | nce: 20, 40, 60 | |
| *dd Settings* | | | |
| Min. AGC Target | | 8.00E3 | |
| Intensity Threshold | | 1.5E5 | |
| Apex Trigger | | -- | |
| Charge Exclusion | | -- | |
| Peptide Match | | Preferred | |
| Exclude Isotope | | On | |
| Dynamic Exclusion | | 10.0 seconds | |

**Table S2. Top metabolites from random forest regression analysis to biomarkers.** Listed are the top metabolites from random forest regression analysis, ranked by increased mean square error (%IncMSE) in decreasing order. If the metabolite was matched on GNPS, the name of the metabolite is listed under the Annotation column. Metabolites with no annotations are designated NM.

| Inflammation | | | | | |
| --- | --- | --- | --- | --- | --- |
| *m/z_*Retention Time | %IncMSE | Cosine Score | Shared Peaks | PPM Error | Annotation |
| X384.234496397944_2.92114275362318 | 0.0224593 |  |  |  | NM |
| X179.062017384554_2.2870659574468 | 0.0146632 | 0.97 | 6 | 1 | Norketamine |
| X465.377877940302_3.03364746376811 | 0.0145534 |  |  |  | NM |
| X547.331989585167_2.98874239130434 | 0.0126831 |  |  |  | NM |
| X401.261138280232_2.26153472222222 | 0.0105595 | 0.95 | 16 | 1 | 7-Oxocholesterol |
| X456.292171168971_2.42917432432432 | 0.0100033 | 0.94 | 12 | 1 | 16:0 Lyso PE |
| X377.325329533161_3.11710398550724 | 0.0093344 | 0.82 | 13 | 7 | Riboflavin |
| X544.339390884811_2.7850829787234 | 0.0092572 |  |  |  | NM |
| X173.078259267423_0.387165277777777 | 0.0091817 | 0.98 | 11 | 2 | Thiamine monophosphate |
| X542.425566903963_2.89093007246376 | 0.0087703 |  |  |  | NM |
| X423.274251937866_2.27051111111111 | 0.0083174 |  |  |  | NM |
| X489.313993883319_2.29389027777777 | 0.0075524 |  |  |  | NM |
| X535.818031642045_2.3068375 | 0.0074338 | 0.77 | 9 | 2 | Ceramide |
| X396.270980140779_2.6986719858156 | 0.0073322 |  |  |  | NM |
| X554.284651472761_2.16796981981982 | 0.0071365 |  |  |  | NM |
| X152.025986676187_2.31203992248061 | 0.0071173 |  |  |  | NM |
| X505.078526973724_0.347101562499999 | 0.0070626 |  |  |  | NM |
| X272.221414082252_3.09807118055555 | 0.0064361 | 0.99 | 8 | 1 | N-Lauroylsarcosine |
| X546.266772101907_2.13186862745098 | 0.0063765 |  |  |  | NM |
| X214.252685003129_0.758197727272727 | 0.0062438 | 0.97 | 4 | 0 | Tetradecylamine |
| X379.248131116231_2.25307361111111 | 0.0062115 |  |  |  | NM |
| X182.081138733267_0.385747222222222 | 0.0061778 | 1 | 7 | 0 | L-Tyrosine |
| X419.273886703863_2.5257069105691 | 0.0057983 |  |  |  | NM |
| X508.041413705955_2.15004224806201 | 0.0056097 |  |  |  | NM |
| X361.747762912657_2.52139634146341 | 0.0054719 |  |  |  | NM |
| X147.076293767056_2.62798333333333 | 0.005379 | 0.99 | 6 | 2 | 2,6-diaminohexanoic acid |
| X185.959649062738_0.247219512195121 | 0.005224 |  |  |  | NM |
| X408.251834450132_2.48886666666666 | 0.0048956 |  |  |  | NM |
| X383.99406135373_0.29435894308943 | 0.0048817 |  |  |  | NM |
| X224.083359890814_2.30365217391304 | 0.004754 | 0.84 | 4 | 1 | Desmethylketamine Isomer |
| X443.445090317144_4.43566097560975 | 0.0047431 | 0.99 | 9 | 2 | C18:1-OH acylcarnitine |
| X110.008889031962_1.53681463414634 | 0.004698 | 0.99 | 5 | 1 | Boc-L-Histidine |
| X254.093790690104_0.914697222222222 | 0.0046553 |  |  |  | NM |
| X348.068694764448_2.39315567375886 | 0.0045604 | 0.96 | 4 | 2 | Inosine 5'-monophosphate |
| X296.18220324634_3.444404964539 | 0.0045408 |  |  |  | NM |
| X282.190500868992_2.12986481481481 | 0.0045185 |  |  |  | NM |
| X522.354968113835_2.86731736111111 | 0.0044836 | 0.97 | 12 | 0 | LPC(18:1) |
| X340.01703202097_0.386235087719298 | 0.0043289 |  |  |  | NM |
| X446.11570072174_2.18165052083333 | 0.0042778 |  |  |  | NM |
| X496.339253039108_2.99043090277777 | 0.0042169 | 0.98 | 12 | 1 | LPC(16:0) |
| X466.980580297009_0.29805 | 0.0041975 |  |  |  | NM |
| X254.093751371016_2.31614148148148 | 0.0040134 |  |  |  | NM |
| X510.257118039014_2.1473906504065 | 0.0040035 | 0.93 | 8 | 1 | LysoPC(17:0/0:0) |
| X331.172034342848_2.19306413043478 | 0.0039609 |  |  |  | NM |
| X547.381310254681_2.78172481481481 | 0.0039266 |  |  |  | NM |
| X435.367163546089_3.16961050724637 | 0.0038956 |  |  |  | NM |
| X137.045746363033_0.48948829787234 | 0.003885 | 0.99 | 7 | 2 | Hypoxanthine |
| X520.33445184727_2.87604097222222 | 0.0038374 |  |  |  | NM |
| X207.056921305295_2.33534964539007 | 0.0038155 | 0.98 | 12 | 0 | Ketamine |
| Fibrosis | | | | | |
| *m/z_*Retention Time | %IncMSE | Cosine Score | Shared Peaks | PPM Error | Annotation |
| X224.083359890814_2.30365217391304 | 0.5808276 | 0.84 | 4 | 1 | Desmethylketamine Isomer |
| X507.327675423971_2.45254227642276 | 0.5185436 |  |  |  | NM |
| X505.078526973724_0.347101562499999 | 0.4628416 |  |  |  | NM |
| X166.086150123951_2.25986423611111 | 0.4211776 | 1 | 7 | 0 | DL-Phenylalanine |
| X662.467313254134_2.64643296296296 | 0.3817735 |  |  |  | NM |
| X690.497773439972_2.75806884057971 | 0.3146294 |  |  |  | NM |
| X701.502526775671_2.74660296296296 | 0.2938911 |  |  |  | NM |
| X669.402624342176_2.4443287037037 | 0.2751121 |  |  |  | NM |
| X649.44879006443_2.93548581560283 | 0.2430406 |  |  |  | NM |
| X403.21146053738_2.51904722222222 | 0.1855925 |  |  |  | NM |
| X348.068694764448_2.39315567375886 | 0.1467101 | 0.96 | 4 | 2 | Inosine 5'-monophosphate |
| X249.115478103609_2.2985152173913 | 0.1379509 |  |  |  | NM |
| X207.056921305295_2.33534964539007 | 0.1255801 | 0.98 | 12 | 0 | Ketamine |
| X536.487689747529_3.84899411764705 | 0.1229489 |  |  |  | NM |
| X799.694066874625_2.82155833333333 | 0.1184275 |  |  |  | NM |
| X660.487912513902_2.74649528985507 | 0.0942147 |  |  |  | NM |
| X391.34096842328_3.2782524822695 | 0.0917223 |  |  |  | NM |
| X390.253105535355_2.47159565217391 | 0.0908728 |  |  |  | NM |
| X721.505637629206_3.02970243055555 | 0.0881036 |  |  |  | NM |
| X440.297135271344_2.56787047619047 | 0.0784888 | 0.99 | 9 | 2 | C18:1-OH acylcarnitine |
| X240.095936883207_2.32600724637681 | 0.0768611 |  |  |  | NM |
| X416.765081405639_2.52563541666666 | 0.072986 | 0.93 | 9 | 2 | C16:0-OH acylcarnitine |
| X493.826529251901_2.46621315789473 | 0.0727893 |  |  |  | NM |
| X230.32451463659_2.78381444444444 | 0.070539 |  |  |  | NM |
| X412.266001259408_2.454181300813 | 0.0693905 |  |  |  | NM |
| X227.222811977284_2.435875 | 0.0690781 |  |  |  | NM |
| X573.544291426495_5.00346097560975 | 0.0680603 |  |  |  | NM |
| X386.238554241417_2.40601493055555 | 0.0678472 |  |  |  | NM |
| X501.375474505106_3.43230104166666 | 0.0676898 |  |  |  | NM |
| X580.513627406529_3.67297809523809 | 0.0643222 |  |  |  | NM |
| X383.13410949707_2.02218632478632 | 0.0634268 |  |  |  | NM |
| X913.606101987161_2.90918958333333 | 0.0632791 |  |  |  | NM |
| X242.075278145926_0.688542857142857 | 0.0627931 |  |  |  | NM |
| X537.484756097561_4.93924390243902 | 0.0617632 | 0.77 | 9 | 2 | Ceramide |
| X716.550028041047_3.22093007246376 | 0.0617376 |  |  |  | NM |
| X272.221462801458_2.82356493055555 | 0.0597848 | 0.99 | 8 | 1 | N-Lauroylsarcosine |
| X781.526081760367_2.9063719858156 | 0.0596803 |  |  |  | NM |
| X511.326920028804_2.30064513888888 | 0.0588435 | 0.93 | 8 | 1 | LysoPC(17:0/0:0) |
| X696.596071139889_2.77282598039215 | 0.0583431 |  |  |  | NM |
| X597.456484587296_3.00889057971014 | 0.0555284 | 0.98 | 7 | 3 | PC916:0/5:0(CHO)) |
| X820.582399370855_3.86742148148148 | 0.0549682 |  |  |  | NM |
| X384.203550985938_2.25196134751773 | 0.0547643 |  |  |  | NM |
| X759.543133824969_2.92635659722222 | 0.0546247 |  |  |  | NM |
| X224.08335543913_2.28234927536231 | 0.0543791 | 0.84 | 4 | 1 | Desmethylketamine Isomer |
| X159.065113248938_0.919428819444444 | 0.0541838 |  |  |  | NM |
| X407.335835000743_2.9747518115942 | 0.0535806 |  |  |  | NM |
| X177.111993156115_3.08455312499999 | 0.0525374 |  |  |  | NM |
| X447.235835594859_2.27897028985507 | 0.0521643 |  |  |  | NM |
| X547.331989585167_2.98874239130434 | 0.0519165 |  |  |  | NM |
| Serum CTGF | | | | | |
| *m/z_*Retention Time | %IncMSE | Cosine Score | Shared Peaks | PPM Error | Annotation |
| X575.412054814996_2.91361956521739 | 59372.544 |  |  |  | NM |
| X386.238554241417_2.40601493055555 | 43187.041 |  |  |  | NM |
| X487.35972151327_3.14305035460992 | 34613.061 |  |  |  | NM |
| X575.411890860787_3.16975905797101 | 15910.46 |  |  |  | NM |
| X331.221451166551_2.47167007575757 | 11899.623 |  |  |  | NM |
| X414.289868644445_5.90893902439024 | 9639.0769 |  |  |  | NM |
| X427.159841025747_2.08864268292682 | 9145.2862 |  |  |  | NM |
| X224.083359890814_2.30365217391304 | 9101.3386 | 0.84 | 4 | 1 | Desmethylketamine Isomer |
| X525.327094293409_2.50704193548387 | 8745.0051 |  |  |  | NM |
| X272.221388524507_2.93454652777777 | 7505.7431 | 0.99 | 8 | 2 | N-Lauroylsarcosine |
| X154.05849543645_0.243820486111111 | 5828.8921 |  |  |  | NM |
| X485.313192972322_2.47777926829268 | 5439.5701 |  |  |  | NM |
| X405.773946810993_2.51673412698412 | 4898.4918 |  |  |  | NM |
| X361.747762912657_2.52139634146341 | 4480.2941 |  |  |  | NM |
| X590.866166277629_2.32253780487804 | 4143.4383 |  |  |  | NM |
| X757.646224189149_2.75794469696969 | 3822.8553 |  |  |  | NM |
| X430.264804280497_2.48171489361702 | 3708.4825 |  |  |  | NM |
| X204.122879046632_0.31105744680851 | 3332.1618 | 0.98 | 6 | 0 | C2:0 acylcarnitine |
| X408.25271621171_2.47855659722222 | 3199.8754 |  |  |  | NM |
| X408.251834450132_2.48886666666666 | 3076.1794 |  |  |  | NM |
| X645.25171866068_2.14965121951219 | 2977.6957 |  |  |  | NM |
| X840.710518680706_2.99745581395348 | 2819.535 |  |  |  | NM |
| X485.331077761766_2.50097967479674 | 2747.3898 |  |  |  | NM |
| X619.437674329731_2.97769548611111 | 2602.3247 |  |  |  | NM |
| X274.175567626953_2.30278492063492 | 2589.234 |  |  |  | NM |
| X707.489967037204_2.98146666666666 | 2503.854 |  |  |  | NM |
| X689.27737505008_2.17057136752136 | 2471.4231 |  |  |  | NM |
| X299.05300558767_2.35741344086021 | 2434.7139 |  |  |  | NM |
| X214.252685003129_0.758197727272727 | 2392.0152 | 0.97 | 4 | 0 | Tetradecylamine |
| X605.422728828761_2.96353333333333 | 2380.9932 |  |  |  | NM |
| X575.412155211566_2.85929528985507 | 2367.8002 |  |  |  | NM |
| X272.086713288959_2.29702631578947 | 2341.0915 |  |  |  | NM |
| X299.998163859207_0.41747738095238 | 2258.1399 |  |  |  | NM |
| X456.292171168971_2.42917432432432 | 2244.4579 | 0.94 | 12 | 1 | 16:0 Lyso PE |
| X575.412390096419_2.97907753623188 | 2119.7012 |  |  |  | NM |
| X454.312859429253_2.65083395061728 | 2117.0903 | 0.93 | 12 | 1 | 16:0 Lyso PE |
| X421.297329711914_5.88518111111111 | 2064.5449 |  |  |  | NM |
| X310.912137167794_0.250892261904761 | 2048.3813 |  |  |  | NM |
| X318.164321473092_2.21703723404255 | 1954.5568 |  |  |  | NM |
| X389.2632528769_2.67620855855855 | 1897.8522 |  |  |  | NM |
| X405.773987205881_2.52675775193798 | 1830.8401 |  |  |  | NM |
| X375.247545567954_2.52281463414634 | 1815.5558 |  |  |  | NM |
| X545.401957636294_3.09573876811594 | 1774.211 |  |  |  | NM |
| X628.497775068115_3.07716773049645 | 1765.6048 |  |  |  | NM |
| X278.539625496699_2.34922413793103 | 1760.7561 |  |  |  | NM |
| X249.115478103609_2.2985152173913 | 1561.4427 | 0.95 | 5 | 1 | Tri(propylene glycol) butyl ether |
| X428.305409749348_5.91522592592592 | 1522.5962 |  |  |  | NM |
| X524.370490447613_3.03744270833333 | 1414.8639 | 0.98 | 10 | 0 | LPC18:0 |
| X502.813790215738_2.30057482269503 | 1351.7758 |  |  |  | NM |
| Serum TGF-β | | | | | |
| *m/z_*Retention Time | %IncMSE | Cosine Score | Shared Peaks | PPM Error | Annotation |
| X701.502526775671_2.74660296296296 | 45834.985 | 0.94 | 7 | 3 | C16 Sphingomyelin |
| X166.086150123951_2.25986423611111 | 16687.774 | 1 | 7 | 19 | DL-Phenylalanine |
| X662.467313254134_2.64643296296296 | 5995.4758 |  |  |  | NM |
| X467.357110391034_2.73451231884058 | 5371.9742 |  |  |  | NM |
| X416.765081405639_2.52563541666666 | 3873.0467 | 0.93 | 9 | 2 | C16:0-OH acylcarnitine |
| X123.055343059457_0.289666319444444 | 2632.3725 |  |  |  | Nicotinamide |
| X822.561354507405_3.63045851851851 | 1921.9599 |  |  |  | NM |
| X778.549647504223_2.74799239130434 | 1881.7288 |  |  |  | NM |
| X207.056921305295_2.33534964539007 | 1872.0719 | 0.98 | 12 | 0 | Ketamine |
| X613.450751428612_2.73802065217391 | 1675.6217 | 0.87 | 25 | 2 | Glutathione |
| X645.440446777343_2.711946 | 1380.5095 |  |  |  | NM |
| X434.279278091531_2.47515775193798 | 1225.3184 |  |  |  | NM |
| X591.385480055176_3.03089255319148 | 1197.4799 | 0.98 | 7 | 3 | PC(16:0/5:0(CHO)) |
| X249.115478103609_2.2985152173913 | 1171.4276 |  |  |  | NM |
| X254.093751371016_2.31614148148148 | 1152.4881 |  |  |  | NM |
| X493.826529251901_2.46621315789473 | 1104.7569 |  |  |  | NM |
| X566.549757794636_4.55284471544715 | 1088.6508 |  |  |  | NM |
| X604.365150237403_2.38967056737588 | 1040.8326 |  |  |  | NM |
| X513.804813957214_2.30130868055555 | 1034.6369 |  |  |  | NM |
| X170.092378144131_2.26928993055555 | 969.1099 | 0.9 | 7 | 0 | 1-Methyl-L-histidine |
| X683.577791050502_4.99589904761904 | 923.6022 |  |  |  | NM |
| X440.297123264622_2.69971936936936 | 898.24403 | 0.99 | 9 | 2 | C18:1-OH acylcarnitine |
| X564.879230499267_2.746175 | 884.63471 |  |  |  | NM |
| X408.25271621171_2.47855659722222 | 869.00823 |  |  |  | NM |
| X575.412155211566_2.85929528985507 | 853.00347 |  |  |  | NM |
| X177.111930586255_3.08792916666666 | 841.26357 | 0.98 | 5 | 0 | Monoethyl phthalate |
| X481.30117285479_2.38662282608695 | 839.77372 | 0.79 | 9 | 1 | LPC(18:0) |
| X394.111005904009_5.79962592592592 | 832.94543 |  |  |  | NM |
| X332.242492382311_2.48492407407407 | 819.26253 | 0.8 | 9 | 0 | C10:0-OH acylcarnitine |
| X254.093760831727_2.18378629629629 | 798.03371 |  |  |  | NM |
| X591.385495275291_3.09514929078014 | 796.51761 | 0.98 | 7 | 3 | PC(16:0/5:0(CHO)) |
| X567.49477410487_5.1715972868217 | 753.69008 |  |  |  | NM |
| X690.497773439972_2.75806884057971 | 746.46589 |  |  |  | NM |
| X346.226934293421_2.45405406504065 | 731.36081 |  |  |  | NM |
| X230.32120164075_2.72050296296296 | 727.54101 |  |  |  | NM |
| X230.170737149594_2.76780222222222 | 727.25209 | 0.93 | 4 | 1 | Isobutyryl-L-carnitine |
| X469.249023379267_2.29397222222222 | 711.77325 |  |  |  | NM |
| X369.296818801734_3.11697872340425 | 705.68645 |  |  |  | NM |
| X718.529602723769_2.77819826388888 | 704.71882 |  |  |  | NM |
| X693.474612862036_2.92641418439716 | 689.20763 |  |  |  | NM |
| X687.486721801757_2.81371333333333 | 676.62823 |  |  |  | NM |
| X607.316024441189_2.52243833333333 | 673.53171 |  |  |  | NM |
| X230.171343239455_2.74933851851851 | 655.79636 | 1 | 7 | 2 | N,N-Dimethyldodecylamine N-oxide |
| X348.721476995126_2.91391775362318 | 634.44378 |  |  |  | NM |
| X449.800222257288_2.52332398373983 | 608.70605 |  |  |  | NM |
| X423.330826893591_2.81694492753623 | 603.57904 |  |  |  | NM |
| X316.211158261602_2.18024131944444 | 598.37479 |  |  |  | NM |
| X799.694066874625_2.82155833333333 | 595.06185 |  |  |  | NM |
| X384.234496397944_2.92114275362318 | 587.34737 |  |  |  | NM |
| Serum PDGF | | | | | |
| *m/z_*Retention Time | %IncMSE | Cosine Score | Shared Peaks | PPM Error | Annotation |
| X666.586415608724_3.78865111111111 | 555700.8 |  |  |  | NM |
| X89.0613639831543_3.19536666666666 | 75218.707 |  |  |  | NM |
| X451.006614140101_0.294282738095238 | 60889.28 |  |  |  | NM |
| X363.30979755283_0.971782575757575 | 45877.871 |  |  |  | NM |
| X298.185454261822_0.761343181818182 | 39334.703 |  |  |  | NM |
| X227.078832081386_1.25716488095238 | 24638.468 | 0.74 | 7 | 2 | L-Carnosine |
| X491.791550006013_2.29344375 | 16523.102 |  |  |  | NM |
| X463.300427246093_2.463456 | 16469.433 |  |  |  | NM |
| X384.114374014047_0.614198717948717 | 16225.102 |  |  |  | NM |
| X828.514923133953_2.26939895833333 | 15760.385 |  |  |  | NM |
| X830.567460885632_3.6424536231884 | 15505.454 | 0.98 | 6 | 2 | PC(20:4/20:4) |
| X723.485258692333_2.84474492753623 | 15311.824 |  |  |  | NM |
| X597.456484587296_3.00889057971014 | 14233.127 |  |  |  | NM |
| X388.304597060592_2.64789037037037 | 12974.992 |  |  |  | NM |
| X502.813790215738_2.30057482269503 | 12261.975 |  |  |  | NM |
| X531.38648889232_0.248 | 11841.001 |  |  |  | NM |
| X429.024539815968_0.362487931034482 | 11482.71 |  |  |  | NM |
| X709.469444402322_2.80366376811594 | 10640.972 |  |  |  | NM |
| X425.752062479654_2.26979861111111 | 9440.7732 | 0.76 | 4 | 38 | 6.beta.-Hydroxymedroxyprogesterone 17-acetate |
| X585.287194228753_2.20268008130081 | 8620.3744 |  |  |  | NM |
| X381.265691245474_2.75893699186991 | 8228.1872 |  |  |  | NM |
| X542.365909576416_2.69998680555555 | 7838.8589 |  |  |  | NM |
| X547.331989585167_2.98874239130434 | 7702.7724 |  |  |  | NM |
| X635.433171364917_2.77170185185185 | 7609.8322 |  |  |  | NM |
| X722.648765563964_5.02950833333333 | 7176.2347 |  |  |  | NM |
| X447.765221595764_2.27704652777777 | 7137.6215 |  |  |  | NM |
| X103.05450711942_0.585831914893617 | 6437.5532 |  |  |  | NM |
| X292.605759811401_1.98244541666666 | 6329.955 |  |  |  | NM |
| X254.0751247406_2.11308452380952 | 6076.9929 |  |  |  | NM |
| X524.370679081758_3.45433333333333 | 5553.8701 | 0.98 | 10 | 0 | LPC(18:0) |
| X447.235835594859_2.27897028985507 | 5533.1303 |  |  |  | NM |
| X567.494841024394_5.22144186046511 | 4869.3595 |  |  |  | NM |
| X549.317575836181_2.31428833333333 | 4840.5609 |  |  |  | NM |
| X707.490355364406_3.24047318840579 | 4570.4429 |  |  |  | NM |
| X577.865033123944_2.50468828828828 | 4536.9681 |  |  |  | NM |
| X230.173598208968_2.77993645833333 | 4456.9647 | 1 | 7 | 2 | N,N-Dimethyldodecylamine N-oxide |
| X702.534387711422_2.98946631205673 | 4376.6129 | 0.94 | 7 | 3 | C16 Sphingomyelin |
| X115.039112574589_1.87350381944444 | 4359.856 |  |  |  | NM |
| X713.428908216238_2.45390634920634 | 4227.191 |  |  |  | NM |
| X386.238751283368_2.86816174242424 | 4153.6779 |  |  |  | NM |
| X489.339045512337_2.71788333333333 | 4124.0153 |  |  |  | NM |
| X540.446578391348_3.09761879432624 | 4083.1044 |  |  |  | NM |
| X73.5304164886474_2.32606 | 4077.1462 |  |  |  | NM |
| X786.162439165791_2.15605797101449 | 4051.0325 | 0.91 | 19 | 3 | Flavin Adenine Dinucleotide |
| X513.804813957214_2.30130868055555 | 3975.3992 |  |  |  | NM |
| X451.214192175288_2.15706979166666 | 3841.089 |  |  |  | NM |
| X286.528385417099_2.31800396825396 | 3802.3236 |  |  |  | NM |
| X241.099227717471_2.37985942028985 | 3778.9157 | 0.73 | 7 | 1 | Tetradecanedioic acid |
| X437.346471079712_2.77967246376811 | 3749.0914 | 0.99 | 9 | 2 | C18:1-OH acylcarnitine |
