## Supplemental Figure for "Alterations to the cardiac metabolome induced by chronic *T. cruzi* infection relate to the degree of cardiac pathology"

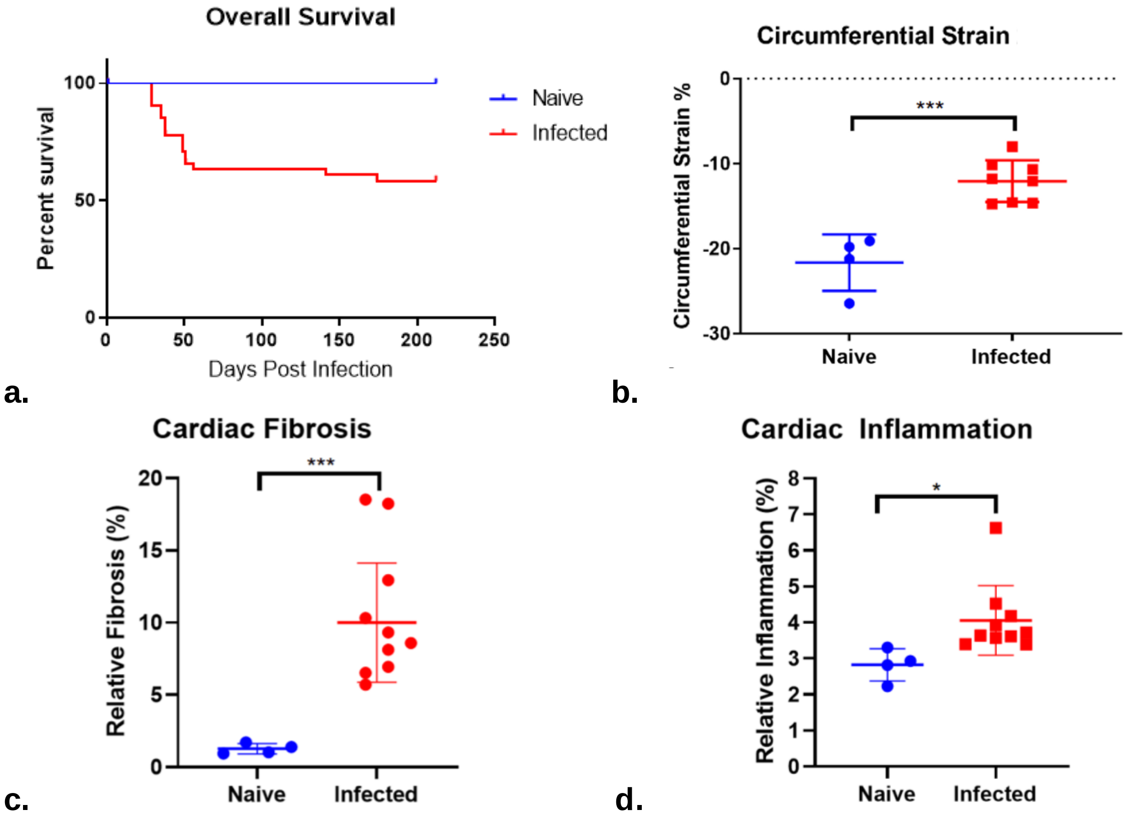


**Figure S1. Survival and cardiac pathology in a mouse model of CCC.** Mortality found to be 40% in the infected group and 0% in the naïve uninfected control group (a). Mortality in the acute phase of the infection was found to be 35% and mortality in the chronic phase was found to be 5% (a). Circumferential strain analysis of short axis echocardiography imaging significantly and positively correlated to cardiac fibrosis measured on histopathology, modified from Hoffman *et al* 2019. (b). Cardiac fibrosis (c) and cardiac inflammation (d) were significantly elevated in infected mice compared to naïve control mice. *p<0.05, ***p<0.001.

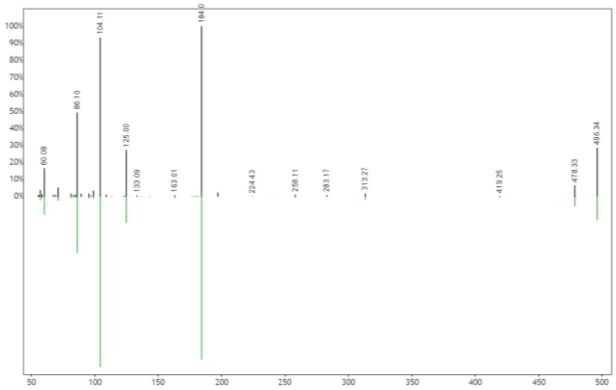

a. LPC(16:0)

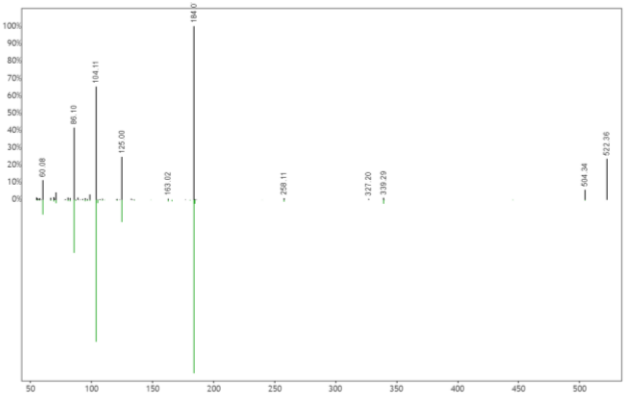

b. LPC(18:1)

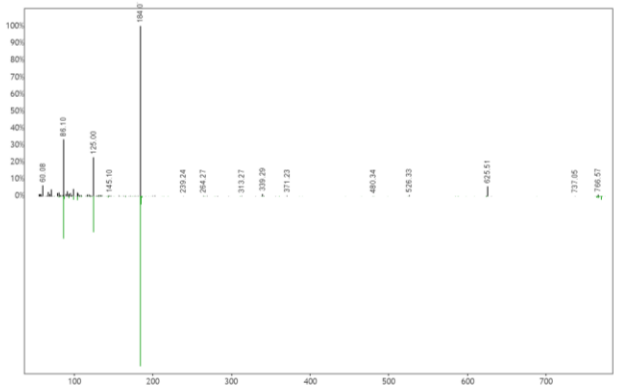

c. PC(o36:5)

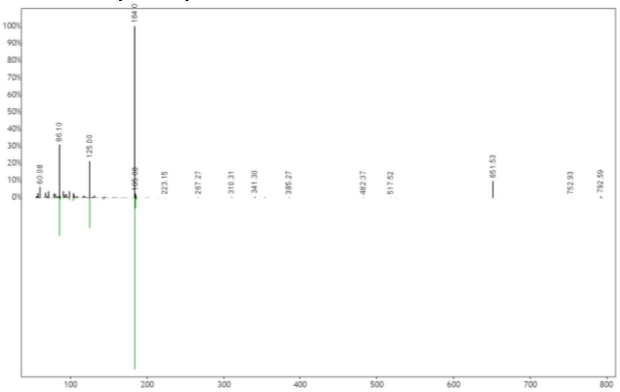

d. PC(o38:6)

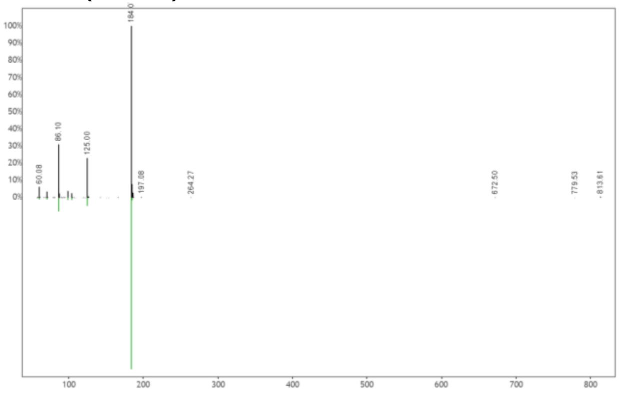

e. C24:1 Sphingomyelin

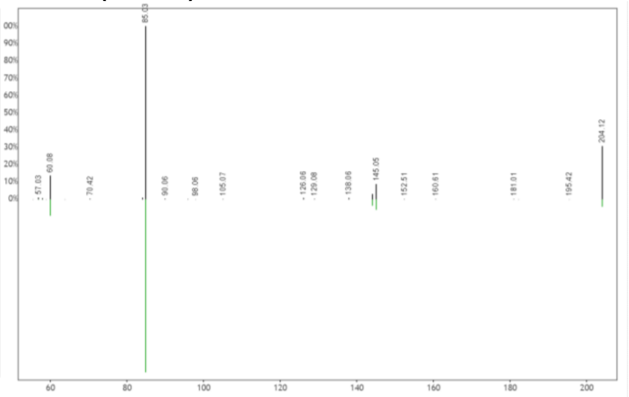

f. C2:0 acylcarnitine

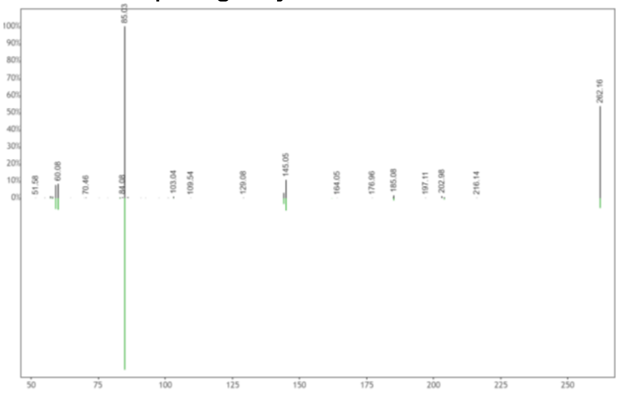

g. C5:0-OH acylcarnitine

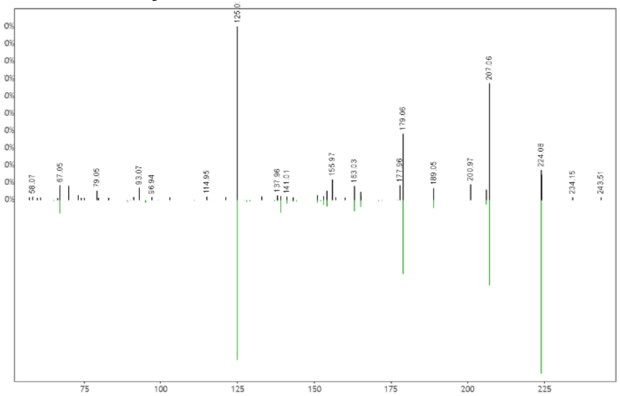

h. Desmethylketamine isomer

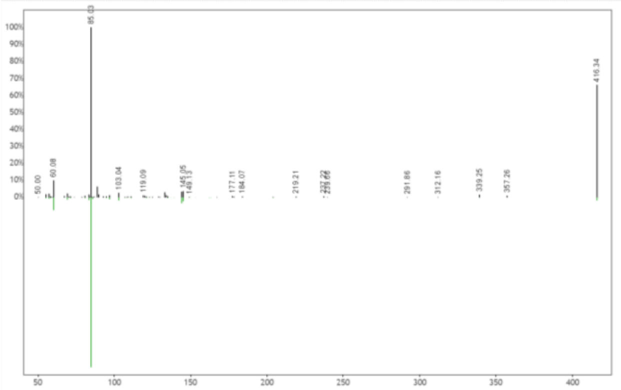

i. C16:0-OH acylcarnitine

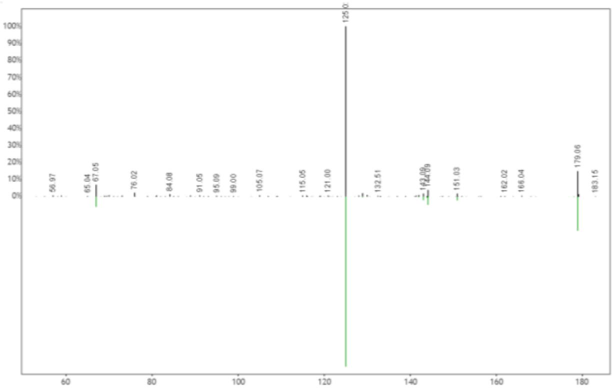

j. Norketamine isomer

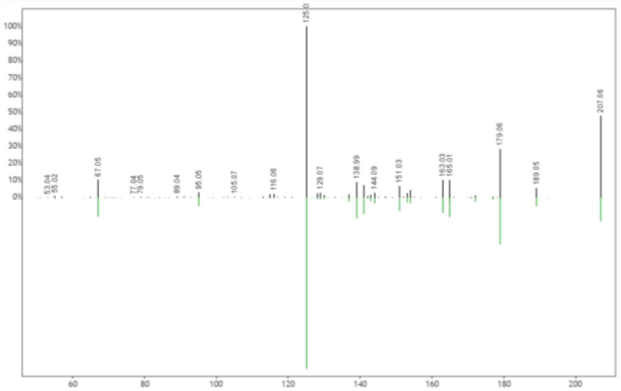

k. Ketamine

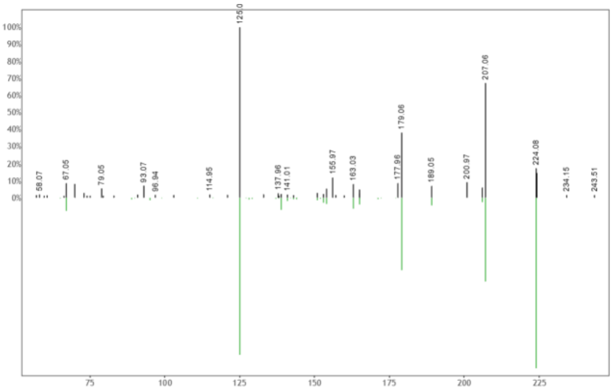

l. Desmethylketamine isomer

**Figure S2. GNPS mirror plots of annotated metabolites.** (a) mirror plot of *m/z* 496.339, RT 2.99s (top, black) to reference library spectrum (LPC(16:0), bottom, green). (b) mirror plot of *m/z* 522.355, RT 2.87s (top, black) to reference library spectrum (LPC(18:1), bottom, green). (c) mirror plot of *m/z* 766.572, RT 5.41s (top, black) to reference library spectrum (PC(o36:5), bottom, green). (d) mirror plot of *m/z* 792.587, RT 5.53s (top, black) to reference library spectrum (PC(o38:6), bottom, green). (e) mirror plot of *m/z* 813.682, RT 3.33s (top, black) to reference library spectrum (C24:1 Sphingomyelin, bottom, green). (f) mirror plot of *m/z* 204.123, RT 0.31s (top, black) to reference library spectrum (C2:0 acylcarnitine, bottom, green). (g) mirror plot of *m/z* 262.164, RT 0.729s (top, black) to reference library spectrum (hydroxyisovaleroylcarnitine, bottom, green). (h) mirror plot of *m/z* 224.083, RT 2.30s (top, black) to reference library spectrum (Desmethylketamine isomer, bottom, green). (i) mirror plot of *m/z* 416.336, RT 2.90s (top, black) to reference library spectrum (C16:0-OH acylcarnitine, bottom, green). (j) mirror plot of *m/z* 179.062, RT 2.29s (top, black) to reference library spectrum (Norketamine isomer, bottom, green). (k) mirror plot of *m/z* 207.057, RT 2.33s (top, black) to reference library spectrum (Ketamine, bottom, green). (l) mirror plot of *m/z* 224.083, RT 2.28s (top, black) to reference library spectrum (Desmethylketamine isomer, bottom, green).
